# Meet me in the middle: median temperatures impact cyanobacteria and photoautotrophy in eruptive Yellowstone hot springs

**DOI:** 10.1101/2021.12.06.471526

**Authors:** Trinity L. Hamilton, Jeff Havig

## Abstract

Geographic isolation can be a main driver of microbial evolution in hot springs while temperature plays a role on local scales. For example, cyanobacteria, particularly high temperature *Synechococcus* spp., have undergone ecological diversification along temperature gradients in hot spring outflow channels. While water flow, and thus temperature, is largely stable in many hot springs, flow can vary in geysing/eruptive hot springs resulting in large temperature fluctuations (sometimes more than 40°C). However, the role of large temperature fluctuations in driving diversification of cyanobacteria in eruptive hot springs has not been explored. Here, we examined phototroph community composition and potential photoautotrophic activity in two alkaline eruptive hot springs with similar geochemistry in the Lower Geyser Basin in Yellowstone National Park, WY. We observed distinct cyanobacterial amplicon sequencing variants (ASVs) consistent with allopatry and levels of light-dependent inorganic carbon uptake rates similar to other hot springs, despite large temperature fluctuations. Our data suggests median temperatures may drive phototroph fitness in eruptive hot springs while future studies are necessary to determine the evolutionary consequences of thriving under continuously fluctuating temperatures. We propose that large temperature swings in eruptive hot springs offer unique environments to examine the role of allopatry vs. physical and chemical characteristics of ecosystems in driving cyanobacteria evolution and add to the debate regarding the ecology of thermal adaptation and the potential for narrowing niche breadth with increasing temperature.

**Importance:** Hot spring cyanobacteria have long been model systems for examining ecological diversification as well as characterizing microbial adaptation and evolution to extreme environments. These studies have reported cyanobacterial diversification in hot spring outflow channels that can be defined by distinct temperature ranges. Our study builds on these previous studies by examining cyanobacteria in geysing hot springs. Geysing hot springs result in outflow channel that experience regular and large temperature fluctuations. While community composition is similar between geysing and nongeysing hot spring outflow channels, our data suggests median, rather than high temperature, drive the fitness of cyanobacteria in geysing hot springs. We propose that large temperature swings may result in patterns of ecological diversification that are distinct from more stable outflows.

## Main Body

Cyanobacteria in hot springs tend to form geographically isolated populations^1,2^ while outflow channel temperature gradients can select for highly adapted, ecologically distinct populations (ecotypes)^1,2^. For example, *Synechococcus* ecotypes are structured by temperature along the stable flow outflow channels of Mushroom and Octopus Springs in the Lower Geyser Basin (LGB) of Yellowstone National Park (YNP), WY, USA^3-8^. In contrast, geysing hot spring outflow channels undergo large temperature fluctuations due to eruptive cycles: continuous flow, a temperature spike from an acute eruption, and a no flow period during source recharge. For the ∼ 500 geysing hot springs in YNP^9,10^, eruption periodicities range from regular (e.g., Old Faithful is ∼ 91-93 minutes) to chaotic (e.g., Steamboat Geyser can vary from 3 days to 50 years^11^). Here we examined phototrophic community composition coupled to rates of light dependent C-assimilation (via ^13^C-labeled bicarbonate microcosms) in the outflow channels of two eruptive hot springs with similar geochemical profiles^12^ (Table S1): Flat Cone (FC) and an unnamed feature we colloquially named ‘The Jolly Jelly’ (JJ; YNP Thermal Feature Inventory ID LFMNN010) in LGB, YNP (Fig 1; Fig S1).

**Figure 1.**
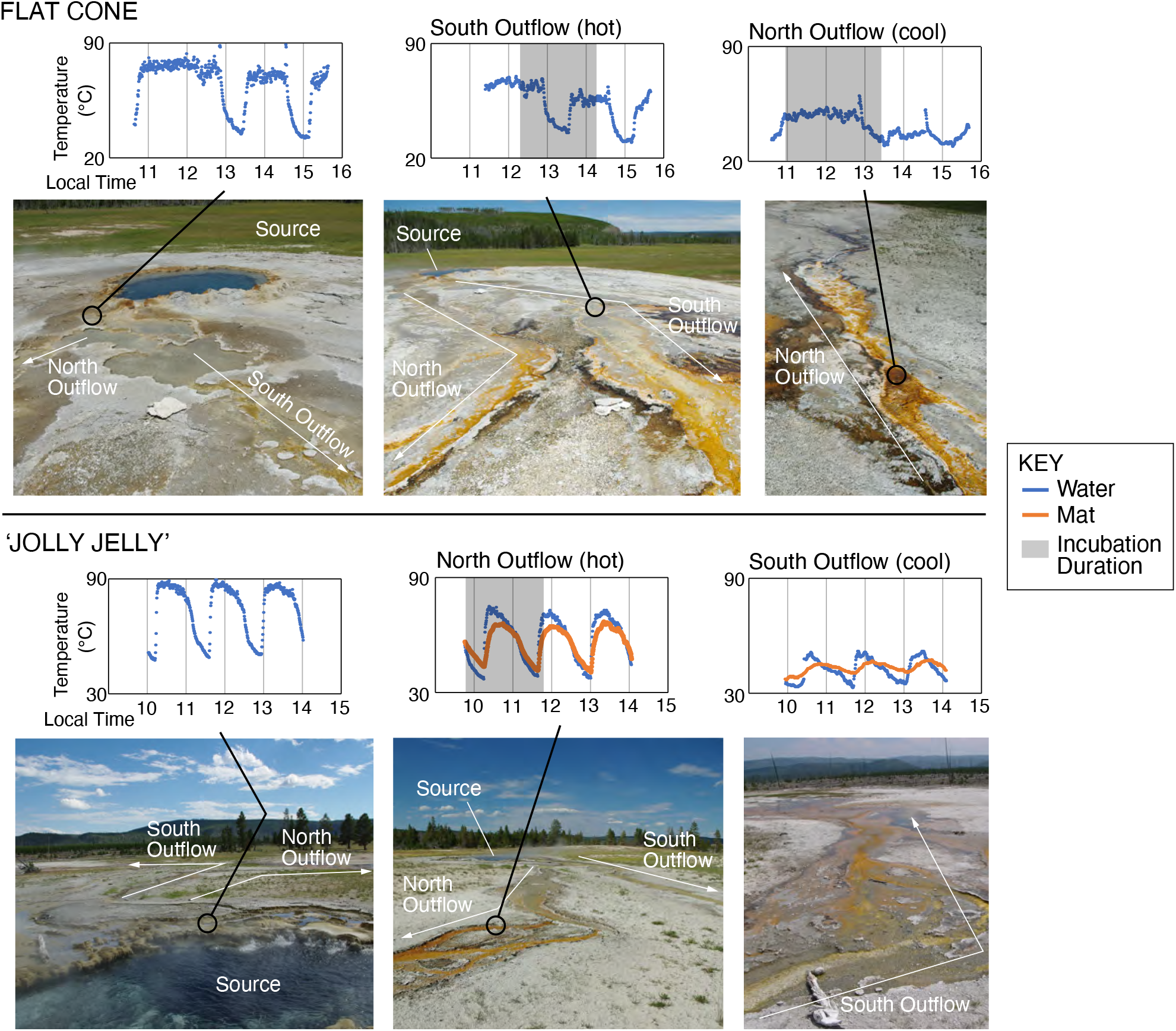
Site photos and temperature variation. Top: temperature measured over a 4-hour window near the FC source and two outflow locations: FC hot (where phototrophs were first visible in the center of the outflow channel) and FC cool. Bottom: temperature measured over a 4-hour window near the JJ source and two outflow locations: JJ hot (where phototrophs were first visible in the center of the outflow channel) and JJ cool. Both sites are located in the Lower Geyser Basin in YNP (SOM Fig 1). Site information (location and select physical and geochemical measurements are provided in Table S1).

FC exhibits a more chaotic eruption periodicity—106 min average, ranging from 25 min to > 12 h, but maintains a steady temperature/outflow rate ∼ 68% of the time (Figs. 1; S2). JJ exhibits a more regular eruption periodicity—88 min average, ranging from 76 to 103 min^13^, with a continuous but fluctuating discharge ∼ 54% of the time (Figs. 1; S3). At FC, phototrophs were first visible in the center of the south outflow channel ∼ 8 m from the source (hereafter ‘FC hot’). Temperatures at FC hot varied by 40.5ºC during a 4 hour observation period: median = 56.0ºC, with maximum = 70.0ºC and minimum = 29.5ºC (Fig. 1). Downstream from the photosynthetic fringe (∼ 14 m from the source, hereafter ‘FC cool’), water reached a median = 40.0ºC over a 4 hour period (maximum = 60.0ºC, minimum = 29.0ºC). At JJ, phototrophs were first visible in the center of the north outflow channel ∼ 24 m from the source (hereafter ‘JJ hot’). At JJ hot, temperatures varied by 38.0ºC during a 4 hour observation period: median = 61.5ºC, with maximum = 75.0ºC and minimum = 37.0ºC (Fig. 1). Further downstream (∼ 60 m from the source, hereafter ‘JJ cool’), median = 42.5ºC (maximum = 52.0ºC, minimum = 33.0ºC). Temperatures deeper in the phototrophic mats were muted compared to that of the water at the mat-water interface (Fig. 1): at a depth of ∼ 1 cm in the JJ hot mats, median = 58.5ºC, with maximum = 67.5ºC and minimum = 40.5ºC.

Despite temperature fluctuations of up to 40ºC, diversity and the composition of putative phototrophs in the geysing sites were similar to those in non-geysing sites (e.g.^14-16^): richness and diversity were lower in phototrophic mats near the upper temperature limit of photosynthesis (Fig. 2A) and, at the 97% sequence identity (defined as operational taxonomic units (OTUs)), sequences assigned to Chloroflexi (*Roseiflexus* and *Chloroflexus*), Cyanobacteria (*Synechococcus* and *Candidatus* Gloemargarita) and Chlorobi (*Candidatus* Thermochlorobacteriaceae bacterium GBChlB) were abundant. Notably, sequences affiliated with other cyanobacteria including *Candidatus* Gloemargarita, *Geitlerinema* PCC−8501, *Leptolyngbya* FYG, and *Pseudanabaenaceae* were only recovered from the ‘cool’ sites consistent with increasing diversity with decreasing temperature.

**Figure 2.**
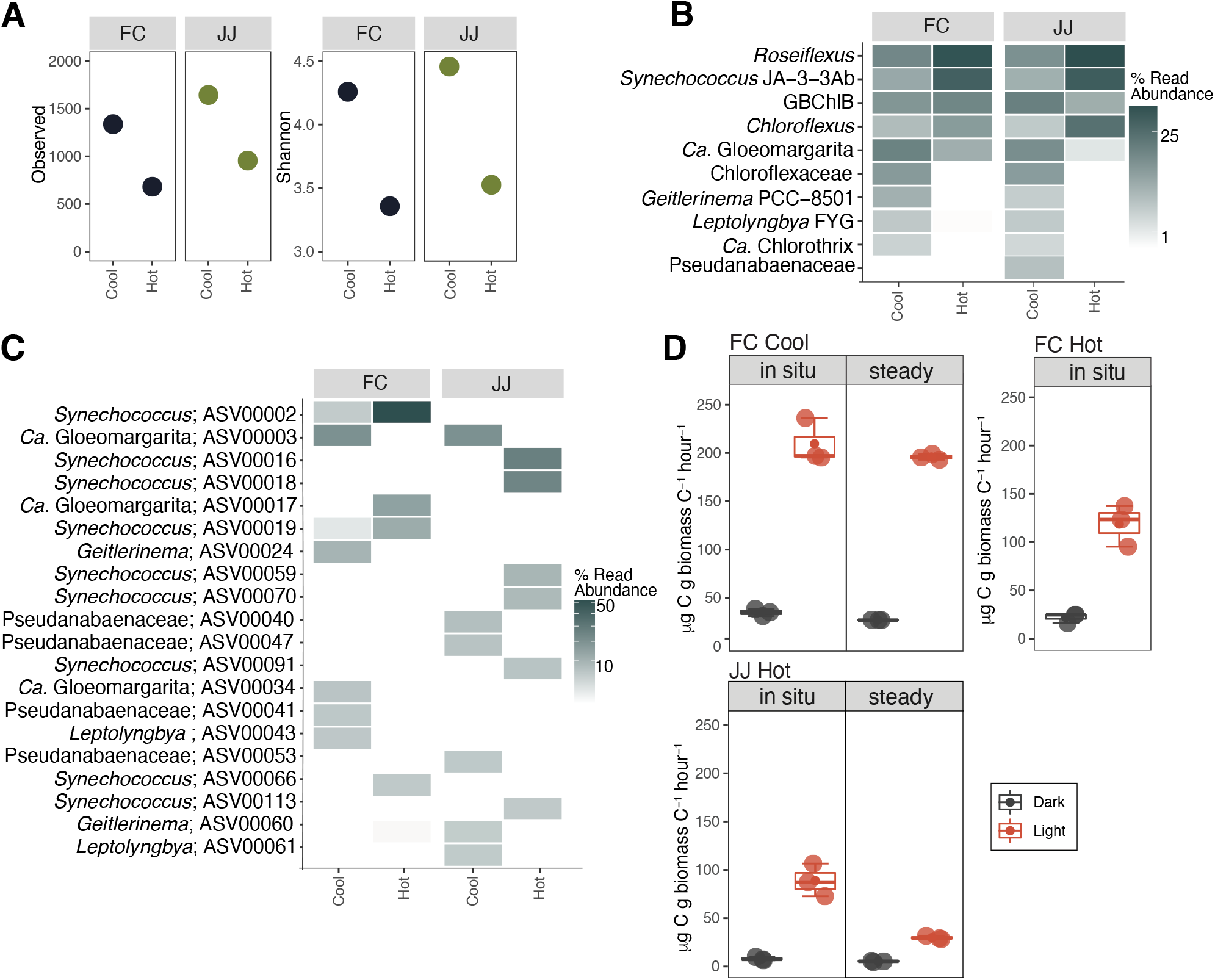
Diversity, phototroph community composition, and C-assimilation rates. (A) Richness and Shannon diversity indices calculated for the 16S rRNA amplicons. (B) Heatmap of the relative abundance of OTUs assigned to putative bacterial phototrophs following Hamilton et al., 2019. (C) Heatmap of the relative abundance of cyanobacterial ASVs. (D) Rates of C-assimilaton in microcosm performed in the dark (wrapped in foil) and light. Error bars from triplicate measurements. In all light vs. dark comparisons, the rates are statistically different (*p* < 0.05). (Rates and *p* values are provided in Table S2). Details of the methods are provided in the SOM.

Temperature selects for distinct cyanobacterial ecotypes in non-geysing outflows (e.g., A’ and A ecotypes occur at higher temperatures while B’ and B are observed at lower temperatures^17^). However, in our geysing outflows, all but one of the most abundant *Synechococcus* cyanobacterial ecotypes (identified as amplicon sequence variants (ASVs)), shared the highest sequence identity with the B’ ecotype. This indicates median temperature (e.g. 56.0ºC at FC hot, 61.5ºC at JJ hot) drives ecotype differentiation in fluctuating systems despite regular exposure to higher temperatures that select for distinct ecotypes in non-geysing systems (e.g. A’ and A ecotypes^17^). With a few exceptions (e.g., ASV00002, ASV00003), the ASVs from JJ and FC were distinct from each other while ASVs from ‘hot’ and ‘cool’ within the same hot spring outflow were also distinct (Fig. 2C). These data are consistent with a role for both geographic isolation and temperature in driving diversification and provide a framework to further examine allopatry vs. physical and chemical characteristics in driving cyanobacterial evolution and diversification under continuously fluctuating temperatures.

We hypothesized that relatively stable temperatures at FC would result in higher rates of photoautotrophy (based on light-dependent C-assimilation rates) compared to JJ, and that the large fluctuation in temperatures at both would result in lower photoautotrophy rates compared to steady temperature sites. We performed microcosms by placing mats and water from hot and cool sites at FC or JJ in sealed serum vials that were amended with NaH^13^CO_3_ following the methods in ^14^. To test our hypotheses vials were incubated: 1) “*in situ*” - vials placed at the sample location, experiencing fluctuating temperatures (see Fig. 1); 2) “steady” - vials placed in nearby non-eruptive hot springs meant to mimic lower temperatures observed at each site (FC cool and JJ hot). As expected, *in situ* rates were higher at FC compared to JJ (Fig. 2D). For *in situ* vs. steady, the C-assimilation rate for the JJ hot mat held at a steady low temperature (steady in Fig. 2D; 28.1 μg C uptake/g C biomass/h) was less than *in situ* microcosms while the C-assimilation rate between *in situ* and steady treatments at FC cool were indistinguishable. Overall, light-dependent C-assimilation rates at both eruptive sites were less than rates observed for alkaline phototrophic communities collected from springs with similar temperate and pH in YNP^14,15^. For example, in previous studies of phototrophic mats, filaments and biofilms from non-geysing alkaline hot springs with similar pH in YNP (e.g. pH 7 -9), observed light-dependent C-assimilation rates ranged from 658.3 to 3813.8 μg C uptake/g C biomass/h^14,15^.

We propose eruptive hot springs are an overlooked but key ecosystem for examining outstanding questions regarding the ecophysiology of hot spring cyanobacteria including does adaptation to increasingly higher temperatures result in narrowing niche breadth^3,18^, what are the roles of temperature and allopatry in driving diversification, and how do Cyanobacteria adapt to high, fluctuating temperatures. Our data indicate stable temperatures might drive higher fitness: light-dependent C-assimilation rates were higher at FC which, while more chaotic in eruption periodicity, supported outflows with stable temperatures 68% of the time compared to more regular eruptivity but continuous temperature variation observed at JJ (changing discharge ∼ 54% of the time). In addition, we recovered sequences most closely related to B’, a lower temperature cyanobacterial ecotype, across a broad niche breadth (at least in terms of temperature). Thus, while median rather than maximum temperature appears to drive cyanobacterial diversification in geysing outflows, the full range of adaptation to high temperature in hot spring *Synechococcus*, particularly in ecotypes from geysing systems, warrants further investigation. Indeed, there is rich history of previous studies on cyanobacterial ecotypes and thus an established comparative framework for examining the evolutionary history and ecophysiology of ecotypes in geysing systems through characterization of new isolates and genomic and metagenomics approaches.

## ACKNOWLEDGEMENTS

T.L.H. and J.R.H. conduct research in Yellowstone National Park under research permit YELL-2019-SCI-7020 issued by the Yellowstone Research Permit Office and reviewed annually. This work was supported by the University of Minnesota. T.L.H was supported by NASA Exobiology award number 80NSSC20K0614. The authors acknowledge the Minnesota Supercomputing Institute (MSI) at the University of Minnesota for providing resources that contributed to the research results reported within this paper. We are grateful to the entire staff of the Yellowstone Research Permit Office for facilitating the permitting process to perform research in YNP. Special thanks to Annie Carlson and Erik Oberg in the Yellowstone Research Permit Office. We thank J. Miller, L. Penrose, L. Brengman, T. Djokic, C. Grettenberger, A. Bennett, A. Rutledge, L. Seyler, and J. Kuether for technical assistance in the field. The authors would like to acknowledge that the research conducted for this work was done in Yellowstone National Park, which was created from land stolen from multiple Native American Nations, especially the Tukudeka (as well as other Shoshone-Bannock and Eastern Shoshone peoples) and Apsáalooke (Crow). These acts were done in part through the guise of Article 2 of the 1868 Fort Bridger Treaty and Article 2 of the 1868 Fort Laramie Treaty. The authors support efforts to give the lands encompassing YNP back to the native peoples who call it home.

## AUTHOR CONTRIBUTIONS

T.L.H. and J.R.H. designed the study, collected samples, and performed the field work. T.L.H. completed the analyses. T.L.H. and J.R.H interpreted the data and wrote the manuscript.

## MATERIALS & CORRESPONDENCE

Correspondence and requests for materials should be addressed to T.L.H.: Trinity L. Hamilton. Department of Plant and Microbial Biology, University of Minnesota, St. Paul, USA, 55108. Phone: +16126256372.

## DATA AVAILABILTY

All sequence data including raw reads with, quality scores for this study have been deposited in the NCBI Sequence Read Archive (SRA) database under with the BioProject number PRJNA756970. Library designations are provided in Table S3.

## SUPPLEMENTAL MATERIALS

## Supplementary Methods

Sample collection and aqueous geochemistry, CO_2_ assimilation (microcosms), nucleic acid extraction and 16S rRNA amplicon sequencing, sequence analysis, and data availability.

**Figure S1**. Sampling locations. Upper Left: Region of the United States that includes the Yellowstone National Park Area. Inset box ‘A’ indicates the region sampled within the Lower Geyser Basin, Yellowstone National Park, USA. Upper Right (A): Western portion of the Lower Geyser Basin including Sentinel Meadows (upper left portion of image) and the Imperial Geyser Basin (bottom center of image). Inset box ‘B’ indicates area of ‘The Jolly Jelly’, inset box ‘C’ indicates area of Flat Cone. Bottom left (B): ‘The Jolly Jelly’. Bottom right (C): Flat Cone. Images provided courtesy of Google Earth.

**Figure S2**. Temperature data logged over time (166 h) near the source of FC in 2008 (Havig 2009).

**Figure S3**. Temperature data logged over time (216 h) near the source of JJ in 2008 (Havig 2009).

**Table S1**. GPS coordinates, pH, conductivity, temperature, and aqueous geochemistry of spring water at the site of sample collection and microcosm incubations. Dissolved inorganic carbon (DIC), DIC δ^13^C values, dissolved organic carbon (DOC), and DOC δ^13^C values are also provided. pH and temperature was recorded at the time of sample collection. Sulf, sulfide; bdl, below detection limit. Detection limits: Nitrate, 0.01 mg/L NO_3-_; Fe^2+^, 20 μg/L.

**Table S2**. Biomass C and N stable isotope analyses of biomass and rates of carbon assimilation and p-values (for each comparison of C assimilation rates). All carbon isotope values are given as absolute values. All p-values are < 0.001. All samples have ^13^C-labeled bicarbonate added. Light, labeled bicarbonate added; Dark, aluminum foil wrapped. St. dev., standard deviation of assays performed in triplicate.

**Table S3**. Accession numbers for the 16S rRNA amplicon libraries included in the present study.

